# pdxBlacklist: Identifying artefactual variants in patient-derived xenograft samples

**DOI:** 10.1101/180752

**Authors:** Max Salm, Sven-Eric Schelhorn, Lee Lancashire, Thomas Grombacher

## Abstract

Patient-derived tumor xenograft (PDX) samples typically represent a mixture of mouse and human tissue. Variant call sets derived from sequencing such samples are commonly contaminated with false positive variants that arise when mouse-derived reads are mapped to the human genome. *pdxBlacklist* is a novel approach designed to rapidly identify these false-positive variants, and thus significantly improve variant call set quality.

**Availability:** pdxBlacklist is freely available on GitHub: https://github.com/MaxSalm/pdxBlacklist

**Supplementary information:** Supplementary data are available.

## 1 Introduction

Patient-derived tumor xenografts (PDXs) are emerging as key preclinical models in oncology research and drug development (Day *et al.*, 2015). PDX models faithfully reflect the molecular characteristics of the source tumour (Tentler *et al.*, 2012) and thus can be used to address diverse subjects such as tumour heterogeneity, evolutionary dynamics and treatment-resistance (Byrne *et al.*, 2017). To this end, major PDX consortia are generating comprehensive molecular profiling data for thousands of models using next generation sequencing (Gao *et al.*, 2015; Bult *et al.*, 2015; Byrne *et al.*, 2017). However, assay sensitivity and specificity is routinely compromised by contaminating mouse DNA and RNA (Lin *et al.*, 2010; Rossello *et al.*, 2013). To limit the technical artifacts caused by sequencing a mixture of two species, each sequence read can be assigned to a source species by comparative alignment to both human and mouse genomes (Conway *et al.*, 2012; Ahdesmäki *et al.*, 2016). However, such read disambiguation imposes a significant computational burden and may be confounded by homologous human-mouse loci (Tso *et al.*, 2014). To complement this strategy and provide an efficient alternative, we developed the *pdxBlacklist* approach for DNA variant calls.

## 2 Approach

*pdxBlacklist* ingests aligned mouse-derived sequencing reads, realigns these to the human genome and outputs a list of artefactual variants resulting from cross-species mapping errors. Once this ‘blacklist’ of *de facto* false positives has been created, it can be used to filter out mouse-related artifacts and refine any PDX variant dataset. Alignment and variant calling are managed by the *bcbio* best-practice variant calling pipelines; this facilitates blacklist generation for diverse variant types (e.g. SNV, indel and SV) and enables genome/aligner/caller combinations to match in-house PDX processing pipelines. The *pdxBlacklist* pipeline is implemented using the *Ruffus* framework (Goodstadt, 2010). For the analysis presented herein, a blacklist was created by processing whole genome sequencing data generated for the Mouse Genomes Project for 18 mouse strains (Keane *et al.*, 2011). Alignment to the hg19 genome build was performed by *BWA-MEM* (Li, 2014), and variants were called using *VarDict* (Lai *et al.*, 2016): this generated a false positive dataset comprising 11,119,424 SNVs, 13,77,355 indels and 2,305,881 complex variants.

## 3 Application

To demonstrate the importance of accounting for mouse-derived artifacts in PDX studies, a *pdxBlacklist*-derived blacklist was employed in three scenarios. First, the blacklist was compared to a public variant database (the COSMIC catalogue (v67) (Forbes *et al.*, 2017)) which identified 41, 675 SNVs that may be PDX artifacts (e.g. COSM1599955 from a glioblastoma PDX study (Yost *et al.*, 2013)). Secondly, the blacklist was used to annotate variants called from 8 synthetic PDX samples that simulate increasing proportions of mouse contamination (Supplementary Figure 1). As expected, the number of blacklist variants detected was strongly correlated with mouse genome contamination (*r*(6)=0.99, *p* = 2.6e-06). Interestingly, a specific variant class was enriched amongst the detected artifacts (CA>TG/TG>CA dinucleotide substitutions). Finally, this blacklist was used to annotate whole exome sequencing derived variant data generated for 300 PDXs, produced using the same analytical pipeline but optionally including read disambiguation by *Disambiguate* (Ahdesmäki *et al.*, 2016). Of the 3,670,468 PDX variants, 83% (3,057,911) are annotated as false positive variants by both *Disambiguate* and *pdxBlacklist*. Despite the considerable redundancy between tools, 122,594 and 12,749 variants are filtered specifically by *Disambiguate* or *pdxBlacklist* respectively (Supplementary Figure 2). Importantly, 111 of the *pdxBlacklist* specific variants are non-synonymous SNVs in Cancer Gene Census members.

## 4 Discussion

*pdxBlacklist* generates false positive variant “blacklists” that can be used to significantly improve confidence in variant calls derived from PDXs. As illustrated, this method is comparable to read disambiguation in sensitivity but offers a significant improvement in performance once a blacklist has been generated. Moreover by using an annotation-based approach, *pdxBlacklist* offers a soft filter rather than a hard filter (as is implicit in *Disambiguate*), and this can be applied to existing PDX variant datasets retrospectively without recourse to the original sequencing reads. Finally, as demonstrated in the final benchmarking exercise, the *pdxBlacklist* output identifies false positive variants masquerading as biologically meaningful variants. To facilitate future use, the blacklist generated for this publication will be included in the *pdxBlacklist* repository. We anticipate *pdxBlacklist* to be an essential tool to serve the ever-expanding community of PDX researchers.

## Acknowledgements

The authors wish to thank Marina Bessarabova and Eike Staub for helpful discussions and comments.

## Author contributions

MS authored the *pdxBlacklist* algorithm and the manuscript. SES & TG generated variant datasets for the PDx cohort.

*Conflict of Interest:* none declared.

